# Hawkmoth flight in the unsteady wakes of flowers

**DOI:** 10.1101/264762

**Authors:** Megan Matthews, Simon Sponberg

**Affiliations:** School of Physics, Georgia Institute of Technology, Atlanta, GA; School of Biological Sciences, Georgia Institute of Technology, Atlanta, GA

**Keywords:** *Manduca sexta*, flight, flower tracking, leading edge vortex, unsteady flow, system identification

## Abstract

Flying animals maneuver and hover through environments where wind gusts and flower wakes produce unsteady flow. Although both flight maneuvers and aerodynamic mechanisms have been studied independently, little is known about how these interact in an environment where flow is already unsteady. Moths forage from flowers by hovering in the flower’s wake. We investigate hawkmoths tracking a 3D-printed robotic flower in a wind tunnel. We visualize the flow in the wake and around the wings and compare tracking performance to previous experiments in a still air flight chamber. Like in still air, moths flying in the flower wake exhibit near perfect tracking at low frequencies where natural flowers move. However, tracking in the flower wake results in a larger overshoot between 2-5 Hz. System identification of flower tracking reveals that moths also display reduced-order dynamics in wind, compared to still air. Smoke visualization of the flower wake shows that the dominant vortex shedding corresponds to the same frequency band as the increased overshoot. Despite these large effects on tracking dynamics in wind, the leading edge vortex (LEV) remains bound to the wing throughout the wingstroke and does not burst. The LEV also maintains the same qualitative structure seen in steady air. Persistence of a stable LEV during decreased flower tracking demonstrates the interplay between hovering and maneuvering.

**Summary statement:** We examined how moths maneuver in the wake of flowers and discover that flower tracking dynamics are simplified compared to still air, while the leading edge vortex does not burst and extends continuously across the wings and thorax.

## Introduction

Flying animals rely on maneuverability to survive in unsteady environments whether evading predators, finding mates, or foraging for food (Dudley 2002*b*, Sprayberry & Daniel 2007, Broadhead et al. 2017). As these animals actively move through their environments, locomotion depends on interactions between their body and the surrounding fluid to produce necessary forces and torques. Changing fluid environments naturally manifest unsteady airflow. Animals must respond to perturbations due to wind gusts and wakes shed from flowers and other objects (Sane 2003, Ravi et al. 2015). Successful flight control requires managing the impact of unsteady flow on body maneuvers and wing aerodynamic forces (Fig. 1). Since biological systems are driven by sensing and feedback, changes to wing forces may also induce body motion and vice versa.

**Figure 1:**
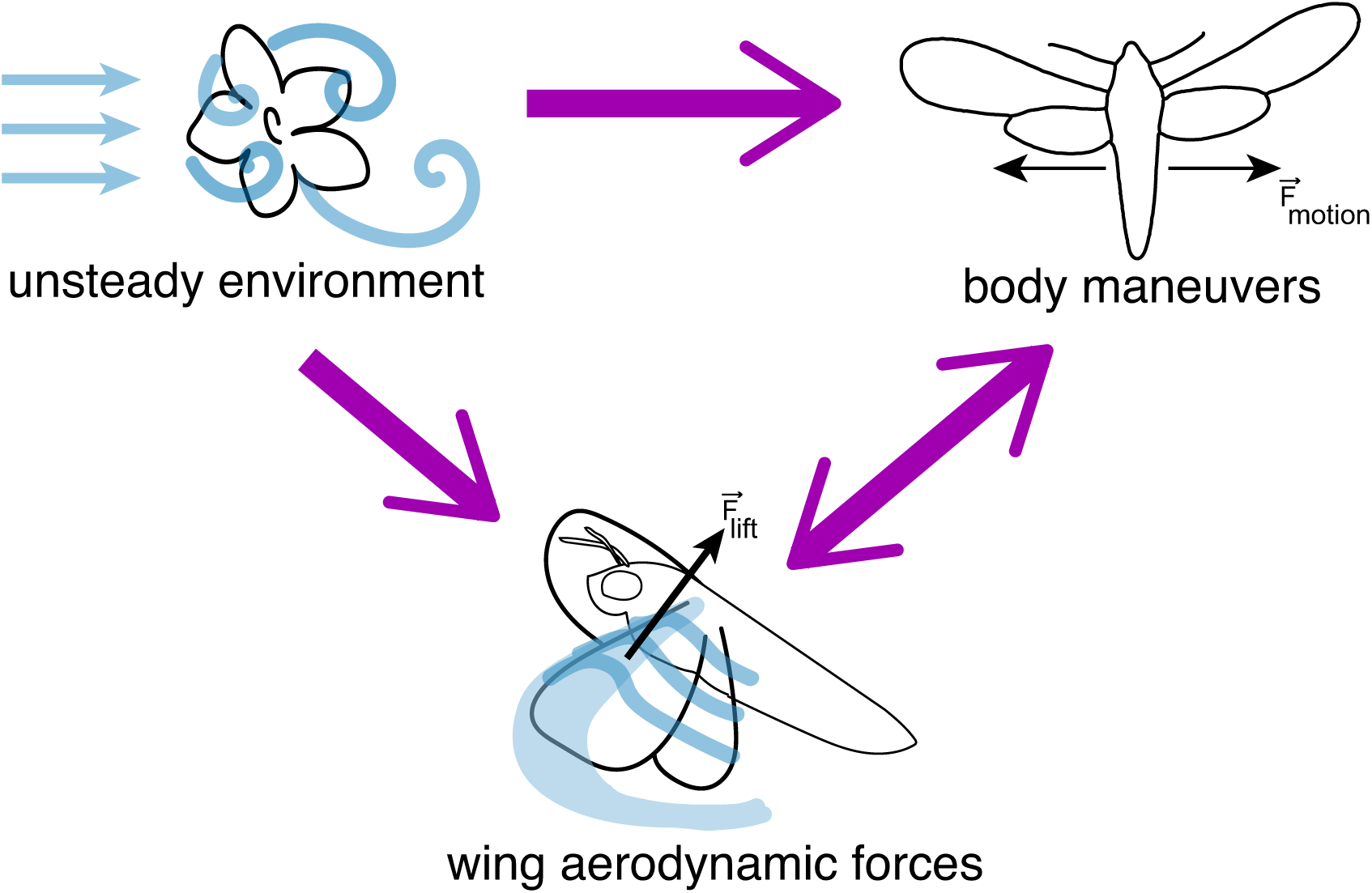
Components for flight success. Insects flying in natural environments must (often simultaneously) (1) interact with unsteady wind, (2) generate stable lift forces through aerodynamic mechanisms, such as the leading edge vortex (LEV), and (3) perform complex body maneuvers to complete tasks. Kinematic variation can arise when the environment pushes the moth or if the moth senses the wake and responds (top arrow). Biological systems are inherently feedback controlled so body maneuvers may also shift if wing aerodynamic forces are changed (double arrow).

How unsteady flow influences flight has been explored for hovering and forward flight (Ortega-Jimenez et al. 2013, Pournazeri et al. 2013, Ortega-Jimenez et al. 2014, Combes & Dudley 2009, Lentink et al. 2010, Ortega-Jimenez et al. 2016, Ravi et al. 2016). However, we still do not know how an unsteady environment impacts the full dynamic range of maneuvers exhibited by flapping fliers. Nor do we know if fundamental aerodynamic mechanisms present in hovering flight persist when an animal must maneuver in flow that is already unsteady. A system identification approach can reveal how flight maneuvers are changed in wind and whether this generalizes across environments (Cowan et al. 2014, Roth et al. 2014).

Hawkmoths must maneuver in unsteady flow while hovering to feed in the wakes of flowers. They must quickly respond to environmental perturbations and changes in flower position (Farina et al. 1995, Sprayberry & Daniel 2007, Sprayberry & Suver 2011, Sponberg et al. 2015, Roth et al. 2016). Using precisely coordinated wing and body kinematics (Sponberg & Daniel 2012), the agile hawkmoth *Manduca sexta*, has adapted robust mechanisms to shift the balance between stability and maneuverability depending on the desired behavior (Dudley 2002*a*, Dyhr et al. 2013). Hawkmoths modulate their kinematics to track flower motion up to 14 Hz, well above what they encounter in nature, albeit with poor performance at high frequencies (Sponberg et al. 2015).

Coordinated wing and body kinematics are also responsible for lift production through unsteady aerodynamic mechanisms (Weis-Fogh 1973, Lighthill 1973). One mechanism, the ubiquitous leading edge vortex (LEV), is thought to contribute to the high lift achieved in insect flight (Ellington et al. 1996, Sane 2003, Dickinson et al. 1999, Chin & Lentink 2016). The LEV has been visualized both qualitatively and quantitatively revealing the basic vortex structure, its dynamics throughout a wingstroke, and vortex stabilization for many insect species (e.g. Srygley & Thomas (2002), Bomphrey et al. (2005, 2009)). However, the LEV on the animal has only been visualized in steady flow (Willmott et al. 1997, Willmott & Ellington 1997*a,c,b*). Moths may need to alter wing motion to maintain lift generation in unsteady flow which could disrupt the quasi-steady LEV. LEV disruption is visualized as vortex bursting, which can occur when increased momentum deflects flow through the vortex core. The deflected flow alters the LEV structure and attachment to the wing (Birch & Dickinson 2001). At the Reynolds numbers for hawkmoth flight, *Re ∼* 10^3^, LEVs on dynamically-scaled flappers burst at high angles of attack (Lentink & Dickinson 2009). However, LEV bursting has not been observed on freely flying or tethered hawkmoths in steady flow (Bomphrey et al. 2005, Johansson et al. 2013, Liu et al. 2018). Although we understand how the LEV generates lift, it is not known how vortex structure and stability are affected by flow in the environment that is already unsteady.

For the hawkmoth, one way unsteady flow is generated in the environment when natural winds encounter flowers. As wind moves around the flower shape, vortices are shed into the wake. Foraging moths feeding from these flowers must interact with these vortices. Feeding in a flower wake introduces two challenges: (1) steady freestream wind and (2) unsteady vortex shedding. In nature, these effects are inseparable and both could have consequences for moth tracking behaviors. The unsteady wake can potentially disrupt tracking maneuvers and the structure of the LEV. LEV bursting and changes to lift production could result in changes to body maneuvers and vice versa (Fig. 1). How do flower wake interactions lead to changes in tracking maneuvers? And how does the LEV interact with unsteady flow already in the environment? In order to address these two questions, we must investigate the interplay between maneuvering, aerodynamics (specifically the LEV), and unsteady flow.

To do this we have moths track robotic flowers in a wind tunnel, where we can control the flow environment. In wind, we also visualize the unsteady wake around the robotic flower, natural flowers, and the moth to reveal impacts on the LEV structure.

If flower tracking performance decreases in wind, then moths must balance reactions to unsteady flow (i.e. the flower wake) with foraging maneuvers. In this case, we predict that moths will react to unsteady flow by matching the dominant vortex shedding frequencies in the flower wake. Consequently, foraging maneuvers used to track the flower will suffer the most at these frequencies.

If the leading edge vortex is disrupted in the flower wake, then we expect to observe LEV growth and bursting around mid-wing during mid-stroke. If no vortex bursting is observed, then we can determine whether LEV structure is altered in the flower wake by visualizing flow over the thorax. The presence of a vortex over the thorax suggests that the LEV is continuous across the full wingspan, rather than conical and rooted on the wings (Bomphrey et al. 2005).

## Methods

### Wind tunnel characteristics

We performed experiments using an open-circuit Eiffel-type wind tunnel (ELD, Inc.). The 150 cm working section consisted of two interchangeable test sections with 60.96 cm symmetric crosssections (schematic drawing in Fig. 2A). The fan is driven by a 3HP induction motor (belt-driven, ODP, 208/230VAC) and generates continuously variable wind speeds from 0.25 - 10 ms^-1^ with less than ±2% variation of the mean. See supplement for specific component details and specifications.

**Figure 2:**
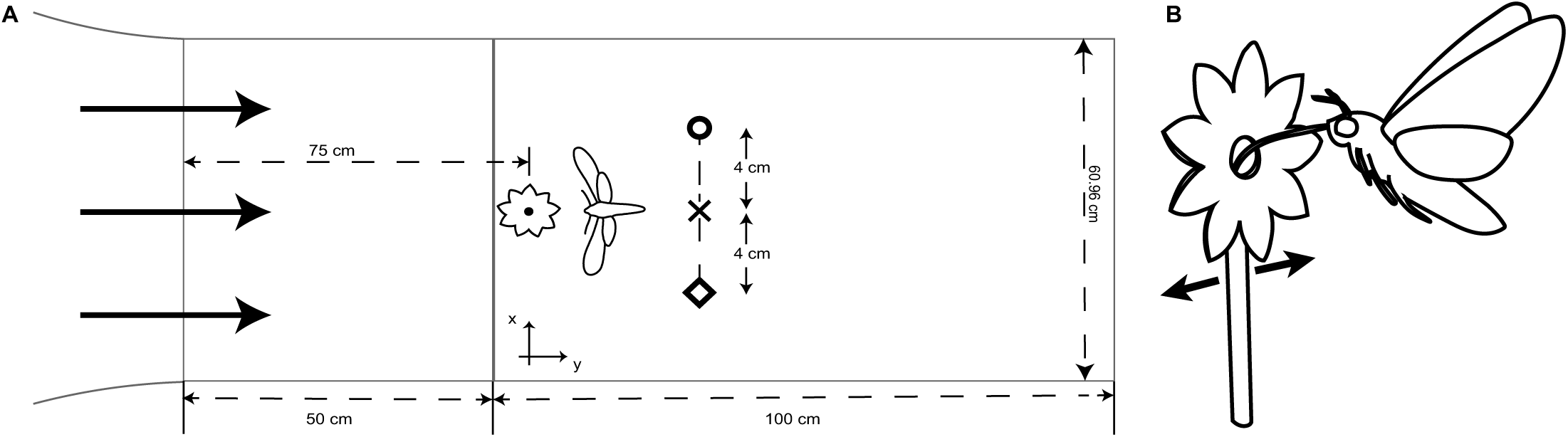
Experimental set-up and conditions. (A) Schematic diagram of the wind tunnel test section. The robotic flower is placed approximately 75 cm downstream (y-direction) of where flow enters and the moth feeds 2-5 cm downstream of the flower. Wind speed measurements were taken at multiple lateral positions (4 cm apart, marked by symbols) with and without the flower (Fig. S1). (B) Sketch of the hawkmoth feeding from the robotic flower. The moth hovers without landing and tracks as the flower laterally oscillates.

### Flow characteristics

We placed a 3D-printed robotic flower (Sponberg et al. 2015) at the front of the secondary test section, 75-80 cm downstream from the upstream mesh and at the approximate lateral midpoint (Fig. 2A, x symbol). The flower is actuated by a bipolar stepper motor (57STH56 NEMA23 with 1067 controller; Phidgets, Inc.) using a 14.5 cm moment arm and the center of the flower face is 20 cm above the bottom panel of the test section.

Wind speed measurements were made at multiple points along the centerline of the working section using an air velocity transducer (TSI Alnor) both with and without the robotic flower present. Measurements were taken at seven downstream points and three lateral positions (Figs. 2A,S1B). The 0.7 ms^-1^ freestream velocity is chosen to replicate what hawkmoths experience in their natural habitat. Anemometer recordings of average wind speeds around six different species of hawkmoth-pollinated flowers all included 0.7 ms^-1^ (Sponberg et al. 2015).

### Experimental set-up and procedure

#### Animals

The hawkmoths used were shipped as pupae from a colony maintained at the University of Washington on a diet that contains retinoic acid (Sponberg et al. 2015). Prior to experiments, the moths were kept on a 12:12 day:night cycle with foraging/feeding time (“dusk”) set around noon EST. Tracking experiments were performed with 5 male and 5 female moths (*m*=1.87 ± 0.54 g, mean ± s.d.) and still flower experiments with 5 male and 5 female moths (*m*=1.70 ± 0.32 g, mean ± s.d.); each moth was only used for a single trial, 2-5 days post-ecolsion, and was not exposed to the artificial flower prior to the experiments. For all trials, naïve moths were dark-adapted to the experiment room for a minimum of 30 minutes. Temperature during experiments was maintained between 24-26°C. A seven-component flower scent (mimicking *Datura wrightii*) is applied to the robotic flower to encourage foraging behavior (Riffell et al. 2014).

Prior to experiments, moths were marked with a dot of white paint (1:1 ratio of tempera and acrylic paint) on the ventral side of the thorax for tracking. Only the thorax point was used since head tracking is not significantly different (Stöckl et al. 2017). Once the moth was feeding (Fig. 2B), we recorded the positions of both the moth and flower throughout the 20 second tracking run. Moths were removed from the flight chamber if feeding was not initiated within 5 minutes.

#### Robotic flower

The robotic flower consists of a 3D-printed flower face (5 cm tip-to-tip diameter) and 2 mL glass nectary attached to the stepper motor. Flower motion is prescribed as a sum of 20 sinusoids (SoS), each with a different driving frequency and randomized phase, and controlled through MATLAB. The stimulus is designed to broadly sample the frequency range and minimize potential learning effects (Roth et al. 2011). A dot of white acrylic paint is applied to the nectary to allow for tracking of the flower motion. The digitized time series verifies that the robotic flower reproduces the designed trajectory (green line, Fig. 3A,B). To limit potential harmonic overlap, the prescribed driving frequencies are logarithmically-spaced prime multiples (0.2-19.9 Hz) (Roth et al. 2011, Dyhr et al. 2013, Roth et al. 2014). To prevent the moths from reaching the saturation limit of their muscle output at the higher frequencies, the velocity amplitude of the flower motion is scaled to be constant at all frequencies (Roth et al. 2014).

**Figure 3:**
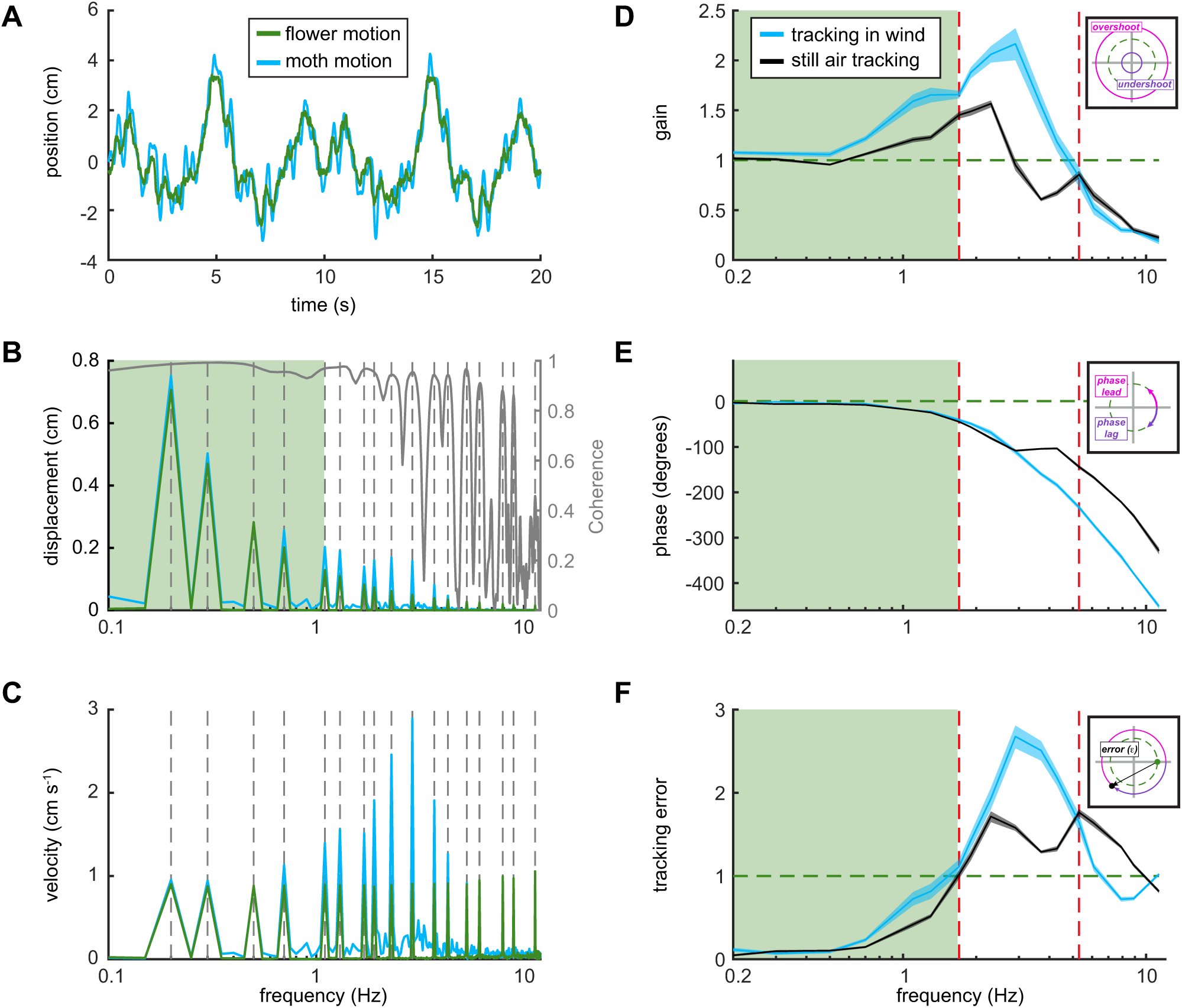
Frequency response comparison for tracking with and without wind. All still air data (black) previously collected in Sponberg et al. (2015). (A) Example raw time series data for one trial of tracking in wind. The moth (blue) overshoots the flower (green) throughout the trial. (B) (left axis) Amplitude (position) in the frequency domain (after Fourier transform of data from A). Peaks correspond to prescribed flower driving frequencies. (right axis) Coherence threshold shows significant tracking drops below 0.9 above 6 Hz. (C) Velocity in the frequency domain. (D,E) Frequency responses (mean ± 95% CI of the mean, two-way ANOVA) for the same tracking task in still (black) and unsteady (blue) air. Responses are categorized into three frequency bands, separated by (red) dashed lines at 1.7 Hz and 5.3 Hz. Gain (D) describes the relative amplitude difference between moth and flower while phase (E) characterizes timing differences. The insets graphically show how gain, phase and tracking error are interpreted in the complex plane. (F) Gain and phase are combined and used to calculate tracking error, the distance from perfect tracking in the complex plane. The green box marks the frequency range below 1.7 Hz matching the range of oscillations exhibited by natural hawkmoth-pollinated flowers.

#### Video recordings

We recorded all flights from below (ventral view) using a Photron UX100 with a 50 mm lens operating at 125 frames per second (fps). Moths were illuminated with two 850 nm IR lights (Larson Electronics) as well as a “moon light” (Neewer CW-126) used to make the flower face visible and set background luminance (Sponberg et al. 2015). Color temperature, based on a blackbody radiation spectrum, was 5400 K. The moon light was equipped with neutral density filters to reduce the measured illuminance to approximately 0.3 ± 0.1 lx (measured in front and to each side of the flower face) for all trials, which is the preferred foraging light level for *M. sexta* (Theobald et al. *2009).*

For smoke-wire visualization an additional Photron UX100 (50 mm lens, 125 fps) was used to record the side view of the animal. The ventral view was used to confirm the horizontal position of the smoke plane relative to either the wingspan or the flower. Additionally, the moon light was increased approximately 0.1 lx to enhance smoke visibility in the ventral camera.

#### Smoke-wire visualization

Smoke visualization (Merzkirch 1987) was performed with a nickel chromium (nichrome) wire aligned with the center of the flower face and approximately 10-20 cm upstream. The 0.25 mm wire was double-coiled and coated with Protosmoke train smoke oil (MTH Trains). After running a current through the wire, the oil condenses into droplets along the length of the wire which are then vaporized into smoke trails as the wire is heated.

### Data analysis

#### Frequency response and tracking performance

After digitizing the flower and moth motions using DLT tracking software (Hedrick 2008), the individual time series (Fig. S5) were detrended and then Fourier transformed to be analyzed in the frequency domain (in MATLAB). Flower tracking has been previously shown to be a linear response (Sponberg et al. 2015, Roth et al. 2016). Using a sum of sinusoids probes a wide dynamic range of behavior, but linearity allows us to generalize to other stimuli. We can characterize the broad frequency response of the system by the gain and phase, (*G, ϕ*), at each unique driving frequency (Fig. 3D,E insets). Details of how gain and phase are calculated are given in the supplement.

#### Tracking error

Instead of separating the real and imaginary components of the frequency response, we can also use the distance from perfect tracking, (1,0), in the complex plane (Fig. 3F, inset) to assess tracking performance.

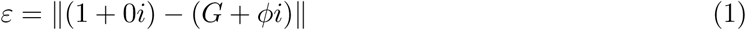

As tracking improves, tracking error approaches zero. Tracking error above 1 indicates that the moth would achieve better performance by remaining stationary at that frequency (Roth et al. 2011, Sponberg et al. 2015).

#### Mechanical (and inertial) power

To assess how body maneuvers specific to flower tracking change due to wind, we restrict our analysis to inertial COM mechanical power. Following previous methods (Sprayberry & Daniel 2007), we calculate the inertial power required to laterally accelerate the center of mass during flower tracking. The moth’s response to the summed sinusoid motion of the robotic flower can be written as

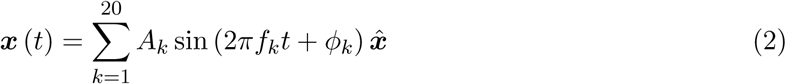

where the indices *k* correspond to the 20 driving frequencies. Lateral oscillations dominate during flower tracking, so we neglect contributions from vertical and looming motion (Fig. S5). Previous experiments with horizontal flower motion also showed that downstream distance from the flower (looming axis) and hovering position (vertical axis) remained fairly constant during tracking, which supports our assumptions (Sprayberry & Daniel 2007). Inertial power (changes in kinetic energy) for the 1D lateral component of tracking motion is given by

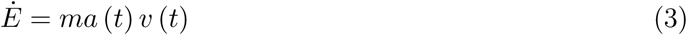

with

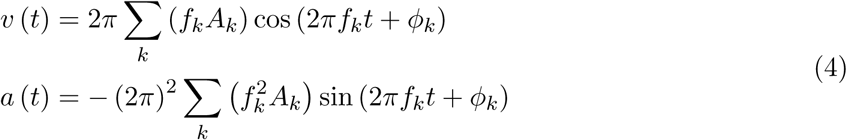

as the lateral components of velocity and acceleration. Then, the inertial power needed for lateral tracking becomes

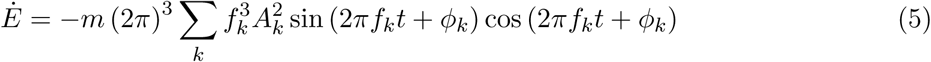

Flower motion is periodic, so the time-averaged tracking power is either positive or negative during each half-period. Since power cycles twice as fast as the underlying driving frequency, we average over a quarter period of flower motion,

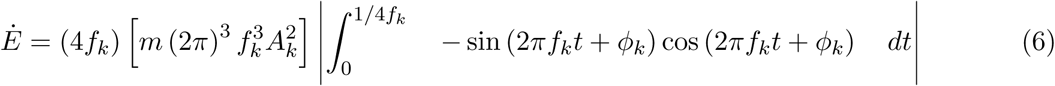

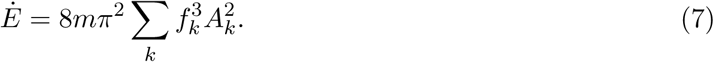

As an upper bound, we calculate inertial power assuming that all changes in kinetic energy must be actively generated. Then, both positive and negative work contribute equally and we take the absolute value to get an upper bound for tracking power (Sprayberry & Daniel 2007). If only positive work must be generated by the animal and energy is dissipated by the environment (e.g. aerodynamic damping (Hedrick et al. 2009)), then the inertial power is exactly half the value in Eqn. 7, which gives a lower bound for inertial power requirements,

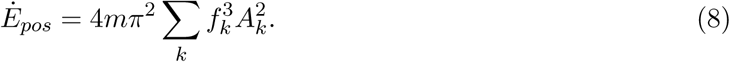

By comparing the inertial COM power for tracking in wind and still air, we can reveal how flower wake interactions affect body maneuvers. Interacting with the unsteady flower wake could alter total mechanical power requirements compared to tracking in still air. Total mechanical power in insect flight combines contributions from multiple sources including profile and induced power, which refer to drag effects on the wings, and parasite and inertial power, corresponding to body drag and acceleration effects (Dudley 2002*a*). While inertial power contains costs for accelerating the wing mass (and added mass), it also includes costs due to accelerations of the body (center of mass, COM) required for flight maneuvers, such as tracking. Body inertial power can be thought of as the added cost to maneuver assuming all other metabolic costs remain constant during maneuvering (e.g. wing inertial power, aerodynamic power, and muscle efficiency (Sprayberry & Daniel 2007)).

#### Statistics

Statistically significant tracking was assessed at each driving frequency using a coherence threshold (Roth et al. 2011, 2014) and Fisher’s exact g test for periodicity (±0.5 Hz frequency bands around each driving frequency were tested with a confidence threshold of 0.05 (Fisher 1929, Sponberg et al. 2015)). Additionally, all averaging and variance estimation were performed in the complex plane (Roth et al. 2011). These values were comparable to results for averaging log gain and using circular statistics to average phase (Roth et al. 2016). Error bars are 95% confidence intervals of the mean unless otherwise noted.

Gain, phase, and tracking error were all tested for significance using two-factor ANOVA (with wind and frequency as our factors). This test was also used to determine significant differences in inertial power within specific frequency bands.

## Results

### Tracking performance decreases in an unsteady flower wake

In unsteady air, all moths successfully tracked for the full 20 seconds with significant tracking at least up to 6.1 Hz with 0.90 coherence (Roth et al. 2011, Sponberg et al. 2015) (Fig. 3B, gray line). Most moths (70%) were able to significantly track up to 11.3 Hz, but few moths (30% or less) were able to track any higher frequencies. This is lower than in still air, where half of the moths significantly tracked flower motion up to 13.7 Hz.

Comparisons between the frequency responses for tracking in wind and still air are best interpreted by examining specific frequency bands: (i) 0.2-1.7 Hz, (ii) 1.7-5.3 Hz, and (iii) 5.3-11.3 Hz. The first band corresponds to the range at which natural hawkmoth-pollinated flowers oscillate (Sponberg et al. 2015, Stöckl et al. 2017), the second captures the range with the largest overshoot in position (Fig. 3B), and the third describes how the moths modulate tracking as they fail.

*(i) Low frequency response: Moths track best at natural flower oscillation frequencies*

The effect of wind on moth dynamics is very small at the lowest frequencies. In wind, moths track nearly perfectly (*G* = 1; *ϕ* = 0) across low frequencies up to 0.7 Hz, revealing only minor differences with tracking in still air (gain and phase difference at 0.7 Hz: 0.15 and 2.8°, Fig. 3D,E). Across all low frequencies, 0.2-1.7 Hz, the moth’s gain was higher in windy conditions than in still air, increasing by 14% at 1.7 Hz (gain for 0.2-1.7 Hz: *F=30.25, dF=1, P<0.05*). However, there were no distinguishable differences in the phase response (0.2-1.7 Hz: *F=0.6, dF=1, P=0.4393*).

Higher gain does not necessarily indicate improved tracking performance. Because the moth was overshooting the flower in windy conditions, the tracking error was larger than in still air (0.7-1.7 Hz, *F=11.14, dF=1, P=0.0012*, Fig. 3F, blue line).

*(ii) Intermediate frequency response: Interaction with wind decreases tracking performance*

As in still air, moths tracking in wind have a distinct region of overshoot (*G >* 1) in the intermediate frequency range (Fig. 3D). However overshoot is both more pronounced and persists over a greater range of frequencies in wind (peak *G* = 2.17 ± 0.16, *ϕ* =109.9°± 5.9°at 2.9 Hz). Overall, tracking in wind between 1.7-5.3 Hz results in a nearly 40% higher peak overshoot (gain: *F=79.52, dF=1, P<0.05*). Tracking in wind also removes the plateau in the phase response, producing a monotonic roll off not seen in still air (phase: *F=5.57, dF=1, P=0.0201*).

Tracking error throughout this frequency band is large due to a combination of gains above 1 and phase lags greater than 90*°*. Maximum tracking error in windy conditions is higher than in still air, with a 70% increase at 2.9 Hz. Tracking error in wind steadily increases until the maximum of 2.68 ± 0.14 at 2.9 Hz (Fig. 3E, blue line) and then decreases until it falls just below the maximum of still air tracking error (1.65 ± 0.11 at 5.3 Hz), resulting in a statistically significant difference with and without wind (*F=58.62, dF=1, P<0.05*). In both cases, but especially with wind, the moth would track these intermediate frequencies better if it stayed stationary (*G* = 0; *ϕ* = 0; *E* = 1).

*(iii) High frequency response: Moths show similar failure dynamics while tracking with and without wind*

Despite the large overshoot in wind at the mid-range frequencies, moths tracking with and without wind fail similarly as flower motion frequency increases towards the saturation limit of the moths’ flight system.

Although few moths were able to successfully track above 11.3 Hz in wind, the decrease in gain leading up to this frequency is similar to the response in still air, with no significant difference (*F=3.19, dF=1, P=0.0773*). The approximately 90°difference in phase lag between wind and still air grows until exceeding a 100°difference at 11.3 Hz (Fig. 3D,E). Above this frequency the moth lags the flower by a full cycle. The continuous phase roll off at high frequency is likely due to an inescapable delay inherent in all real biological systems. Since tracking error is a distance in the complex plane and phases of 0°and ±360°are equivalent, the increased phase lag for tracking in wind reduces tracking error back toward a value of 1. This results in a maximum difference of 0.63 ± 0.03 between tracking in wind and in still air (*F=36.31, dF=1, P<0.05*).

These differences for tracking with and without wind lead to the differences in tracking error (Fig. 3F). In wind and still air, moths fail by undershooting and lagging behind the flower at higher frequencies until, at the highest frequencies, they are effectively non-responsive (*G* = 0; *ϕ* = 0). As tracking gain approaches zero, the tracking error necessarily approaches 1.

### Flower tracking in wind manifests simpler dynamics

The change in the transfer function (frequency response, Fig. 3D,E and Fig. S2) suggests that windy conditions simplify tracking maneuvers. Tracking in wind can be described by a reduced-order dynamical system, compared to still air. While the low frequency behavior is maintained, the response at high frequencies is diminished. The transfer function describing still air tracking includes a simple delay term (Sponberg et al. 2015, Roth et al. 2016, Stöckl et al. 2017) and a minimum of four poles and three zeroes to capture the double peak in gain, but more importantly, the plateau in phase between 3-5 Hz. In wind, the phase plateau is removed and gain has a single peak, so tracking can be described by a lower-order transfer function with only two poles. This order reduction suggests that wind acts as an environmental filter that modifies tracking dynamics at and above the range of vortex shedding. The second gain peak in still air represents a removed pole between 4.3-6.1 Hz, which overlaps the end of the range of vortex shedding frequencies.

### Dominant vortex shedding frequencies of unsteady flower wakes coincide with specific frequency bands

*(i) Roboflower sheds vortices in the intermediate frequency range, matching the frequency band of overshoot*

To explore the temporal dynamics of the unsteady flow around the moth we imaged the wake of the stationary robotic flower. Using smoke-wire visualization, we observed that the dominant vortex structures in the flower wake are irregular, but mostly within the intermediate frequency band of 1.7 to 5.3 Hz. The vortex shedding frequency was determined by observing the number of vortices (rotating in the same direction) over 300 frames, approximately 5-10 cm downstream of the flower face (Fig. 4A, red box). Averaging across four videos, the vortex shedding frequency from the top petals was 2.16 Hz ± 0.25 Hz (mean ± s.d.). Flow around the nectary shed vortices at 0.95 Hz ± 0.17 Hz (mean ± s.d.). However, this is a lower bound of the vortex shedding frequency because the flower sheds vortices that rotate in multiple directions with one or more arriving at the same downstream location simultaneously (Movie S1).

**Figure 4:**
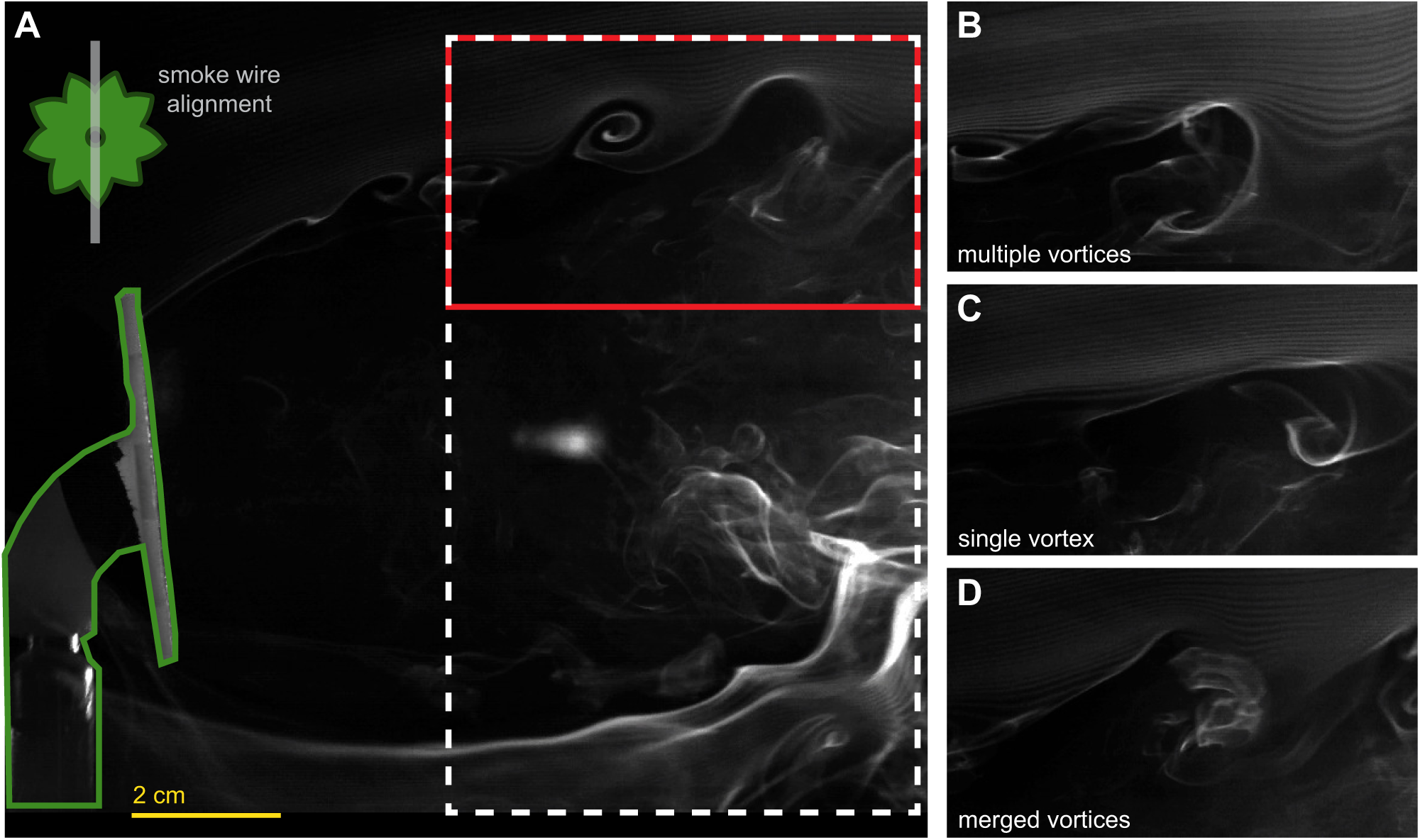
Smoke visualization of the robotic flower wake. (A) Full frame view of flower wake. Inset shows smoke wire lateral alignment with the flower face. The moth primarily feeds in the relatively low flow region approximately 2-5 cm downstream of the flower. Vortices were most distinguishable around 5 cm downstream (white dashed box) and vortex shedding frequency was measured at this location (red box). (B) Snapshot of flower wake (from red boxed region) showing multiple vortices, rotating in multiple directions, passing through the same location. (C) Snapshot of a single vortex. (D) Snapshot of diffuse streaklines due to merging vortices. Level adjustments were made to highlight the smoke lines using Photoshop with a mask over the robotic flower, shown in green.

While the streaklines had a slight upward drift due to higher temperature than surrounding air, the flower wake structure is distinct from the undisturbed streaklines above the flower. As the wake develops downstream the shed vortex structures interact with one another. Some vortices cluster (multiple vortices, Fig. 4B), appear distinct (single vortex, Fig. 4C), or merge with counter-rotating, neighboring vortices (diffuse streaklines, Fig. 4D). These overlapping vortices (Fig. 4B,D) potentially double the number of vortices, raising the estimated dominant vortex shedding frequency up to 4.33 Hz ± 0.49 Hz (mean ± s.d.) from the top petals and 1.90 Hz ± 0.34 Hz (mean ± s.d.) from the nectary. While the moth feeds in a relatively low flow region (approximately 5 cm downstream), the wingspan extends past the face of the flower so structures shed from the perimeter interact directly with both wings. The vortex shedding frequencies range from 2-5 Hz, overlapping the region of increased overshoot (Fig. 3D).

*(ii) Real hawkmoth-pollinated flowers shed unsteady wakes at low frequencies*

Size and material differences between our robotic flower and natural flowers could lead to different unsteady wakes. Hawkmoths forage from flowers of various sizes, from 1-2 cm up to 10 cm (Sprayberry & Suver 2011, Sponberg et al. 2015) and natural flowers are more flexible than the 3D-printed roboflower. Using the same visualization method, we observed the wakes shed from fully bloomed *Datura sp.* (tip-to-tip, flower face diameter: 9 cm, Fig. 5A) and *Petunia sp.* flowers (tip-to-tip, flower face diameter: 7 cm, Fig. 5B) attached to the same rigid support used for the robotic flower. Larger *Datura* flowers shed vortices 2-3 cm further downstream than *Petunia*. This includes the feeding position of the moth (Fig. 5A). Fewer vortices appear in the measurement region (Fig. 5, red box) resulting in a lower frequency of structures shed off the flower petals, (lower bound: 0.43 Hz ± 0.06 Hz; upper bound: 0.87 Hz ± 0.12 Hz). The vortex shedding frequency around the nectary was similar to the roboflower since the same support structure was used (bottom wake structure, Fig. 5A). Averaging over three *Datura* videos gives a lower bound of 0.88 Hz ± 0.26 Hz and an upper bound of 1.77 Hz ± 0.51 Hz. For *Petunia*, these vortices are partially disrupted due to interactions with the lower set of petals, which did not allow for measurement of vortex shedding in this region (Fig. 5B). Based on three different *Petunia*, and averaged over four videos, the vortex shedding frequency is 0.95 Hz ± 0.18 Hz. Unlike the robotic flower wake, *Petunia* showed fewer overlapping vortices, but vortices were still multi-directional, so while an upper bound of 1.90 Hz ± 0.36 Hz is unlikely, some structures are shed at frequencies above 1 Hz.

**Figure 5:**
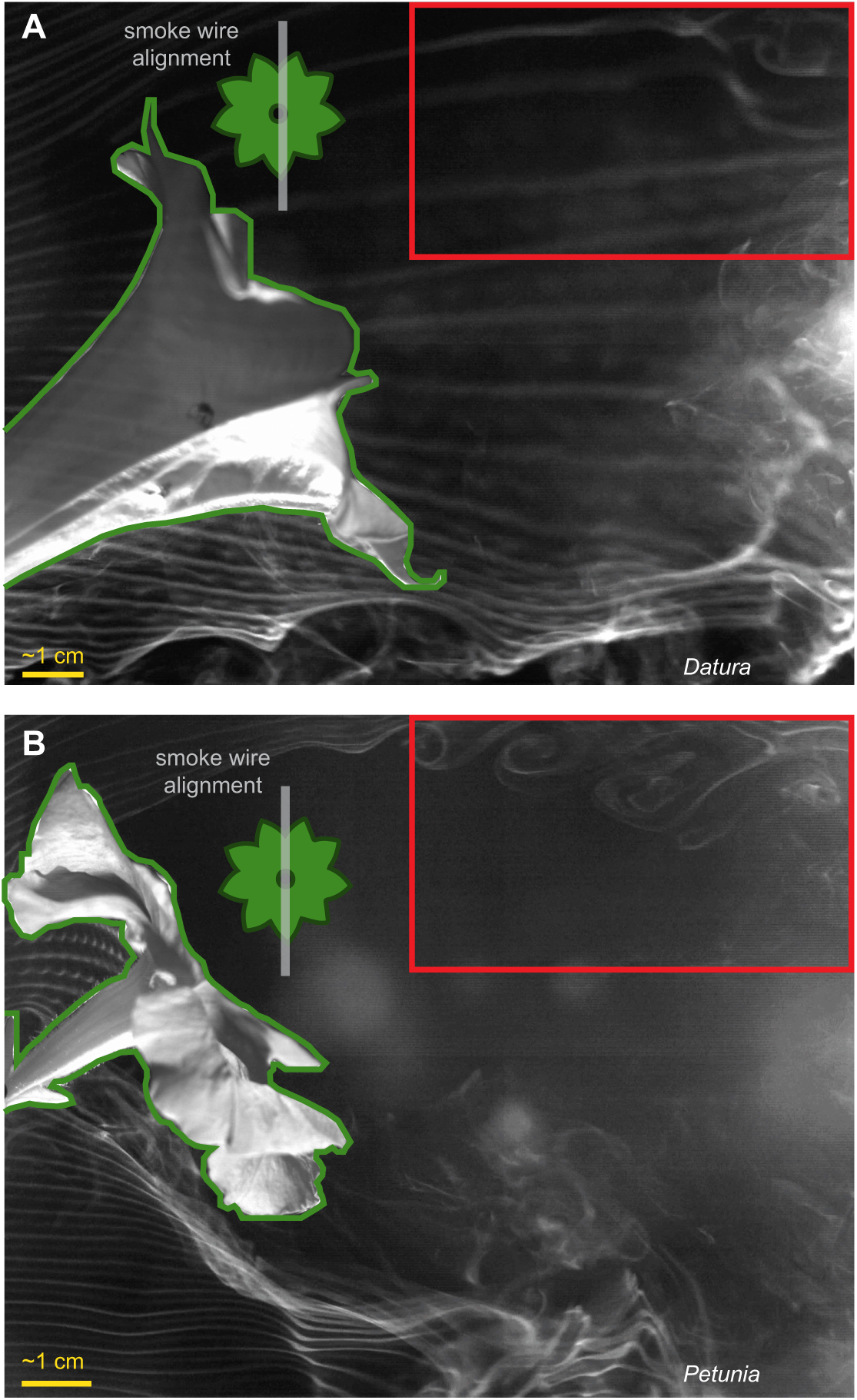
Smoke visualization of natural flower wakes. (A) Snapshot of *Datura sp.* wake. (B) Snapshot of *Petunia sp.* wake. Inset shows smoke wire lateral alignment with the flower face. For both flowers, vortices similar to those shed by the robotic flower are seen coming from the top petals, with fewer passing through the measurement region for *Datura* (red box). The wake structure from the bottom petals is disrupted by the rigid support rod. Global adjustments were made to brightness, contrast, and gamma, within the Photron software (PFV). Additional level adjustments were made to highlight the smoke lines using Photoshop with a mask over the flower, shown in green.

### Inertial power comparisons quantitatively confirm consistency of flower wake and reveal nonlinearity

To test if flower tracking in wind (case 1) is a linear superposition of stationary hovering in wind (case 2, Fig. 6A,B) and tracking in still air (case 3), we compared the mechanical power utilized during these maneuvers. For the two tracking cases, (1) and (3), moths exhibit inertial COM power peaks at each of the driving frequencies and minimal power at the non-driving frequencies (Fig. 6C). When the flower is held stationary in wind (case 2), we expect that power requirements should only increase in the frequency band corresponding to vortex shedding. A stationary flower would result in a stationary moth and therefore low inertial COM power. Then, inertial COM power for hover-feeding in wind (case 2) reveals the maneuvers induced by flower wake interactions (Fig. 6D). Agreement between tracking (case 1) and hover-feeding (case 2) in wind at the non-driving frequencies suggests that the flower motion at various frequencies does not significantly change the vortex shedding frequency. The sum of the moth’s inertial power for hover-feeding in wind (case 2) and tracking in still air (case 3), give a linear prediction for the power needed to track in wind (case 1).

**Figure 6:**
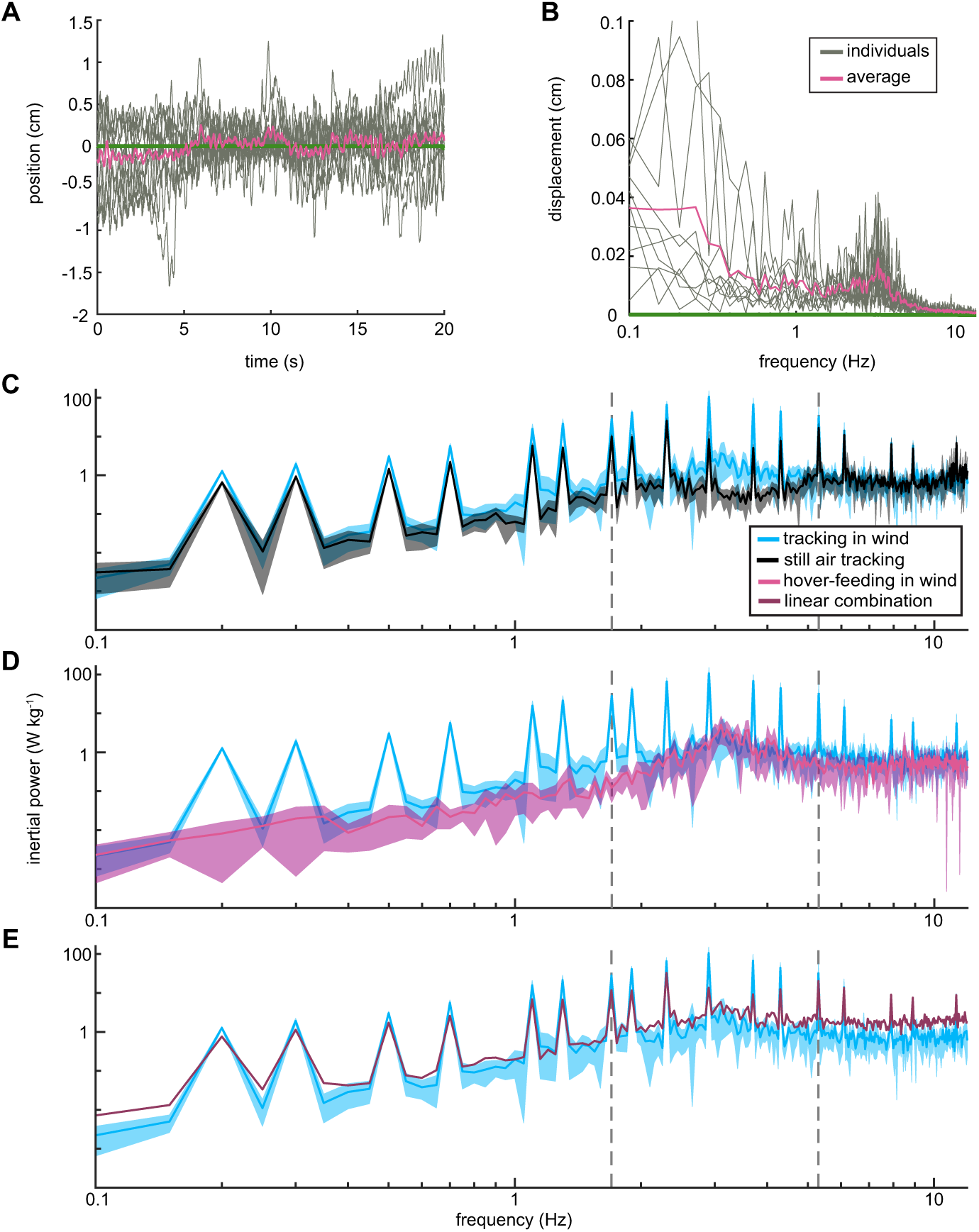
Effects of unsteady wake on inertial power. (A) Example time series data for hover-feeding trials in the wind tunnel. The flower (green) remains stationary while the moth oscillates and tries to maintain a stable position with 0.7 ms^-1^ freestream wind. Traces showing trajectories of all sampled moths (grey) and the mean (pink) show high variation between individuals. (B) Fourier transform of data (from A). Each individual trial was transformed and then averaged. Despite individual variation, all moths display large amplitude oscillations below 1.7 Hz with an additional (smaller) peak occurring between 2-5 Hz. (C) Inertial (COM) power comparison (mean ± 95% CI) for tracking in wind (blue) and in still air (black). Power peaks at the driving frequencies for both tracking cases, but peaks are higher for tracking in wind. (D) Comparison for tracking in wind (blue) and hover-feeding in wind (pink). Power peaks at the driving frequencies for tracking in wind, but both traces show agreement in non-driving frequencies between 1.7-5.3 Hz. (E) Comparison between measured tracking in wind (blue) and the linear combination of still air tracking (Sponberg et al. 2015) and hover-feeding in wind (dark purple). While the linear combination agrees fairly well with the response at the lowest frequencies, it under-predicts the response between 1.7-5.3 Hz and over-predicts at the highest frequencies.

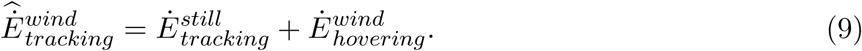

The linear sum 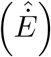 over-predicts the measured response 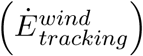 for tracking in wind at all non-driving frequencies (Fig. 6E). The response at the lowest driving frequencies is well-captured by a linear combination of still air tracking and hover-feeding in wind. However, at the mid-range driving frequencies (1.7-5.3 Hz), the linear sum consistently under-predicts the actual response by a minimum of 11.4 W kg^-1^ (at 5.3 Hz) and a maximum of 92.1 W kg^-1^ (at 2.9 Hz). Although there is a slight increase in inertial power between hover-feeding (case 2) and tracking in wind (case 3), the over-prediction by the linear sum, especially between 2-5 Hz (Fig. 6E), is too large to be explained by differences in flower wake characteristics alone.

### Key features of the leading edge vortex are maintained in wind

Since the flower wake significantly decreases tracking performance, we next used smoke visualization to see if the leading edge vortex bursts in wind. Bursting is expected to occur along mid-span during the middle of the wingstroke. In the absence of bursting, the LEV may maintain the same structure observed in steady air with a relatively constant diameter extending across the full wingspan during mid-wingstroke (Bomphrey et al. 2005).

With the smoke wire aligned at mid-wing, we observe a single LEV that reattaches without bursting (Fig. 7A), consistent with LEV structure in steady air conditions (Ellington et al. 1996, Bomphrey et al. 2005). During each downstroke, the mid-wing LEV grows until it is shed prior to the beginning of the upstroke. The stable LEV is most visible at mid-downstroke (Fig. 7A and Movie S2). Although the freestream velocity in our experiments was slightly lower than in previous studies (Ellington et al. 1996, Bomphrey et al. 2005), the mid-wing LEV is qualitatively similar in size (relative to the wing chord) and shape to their results. Other features of the LEV structure, such as trailing-edge (Fig. 7B, white arrow) and tip vortices (Fig. 7A, yellow arrow) are also visible on some wingstrokes, but the full vortex loop structure cannot be resolved with smoke-wire visualization alone.

**Figure 7:**
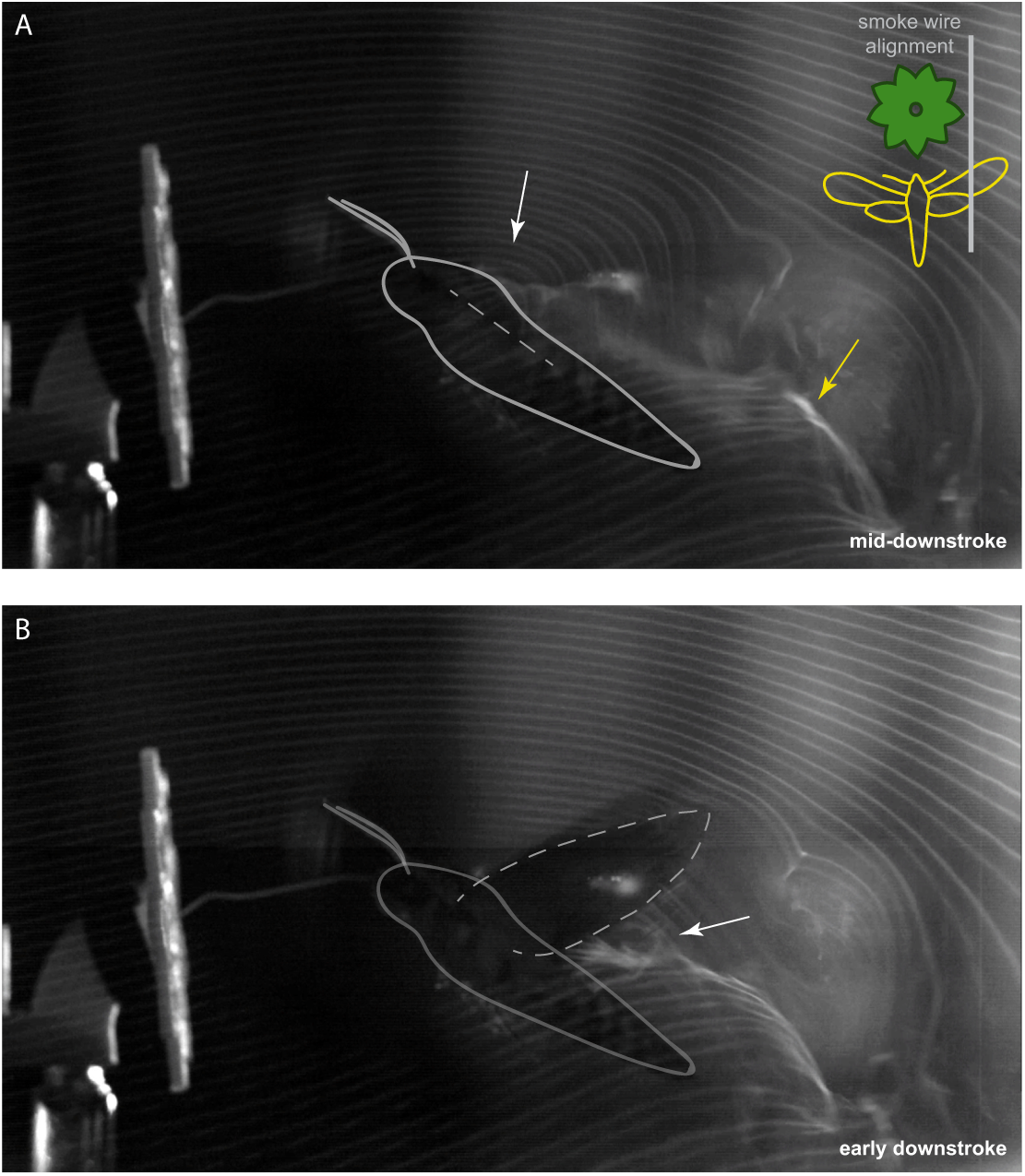
Smoke visualization of the leading edge vortex at the mid-wing position. Free-flying moths (*n* = 2) maintaining a stable position while feeding from the robotic flower in wind. Outline of moth added for clarity. (A) Mid-downstroke. Separated flow region of LEV (white arrow) and roll up of the tip vortex (yellow arrow). The LEV during the mid to late downstroke resembles what has been seen previously for tethered *Manduca* in steady air (compare to Bomphrey et al. (2005)), but the tip vortex (yellow arrow) shows a down and backward trajectory, rather than back and upwards. (B) Early downstroke. A possible trailing-edge vortex (TEV) indicated by a white arrow. Level adjustments were made to highlight the smoke lines using Photoshop. Inset shows smoke wire lateral alignment with the moth and flower face.

The LEV structure is continuous across the thorax in the absence of vortex bursting. Over the thorax, the LEV forms during stroke reversal and grows during the upstroke (Fig. 8B and Movie S3), consistent with observations in steady air (Bomphrey et al. 2005). However, a transient LEV was sometimes present during the downstroke (Fig. 8A), so LEV structure may not be conserved from wingbeat to wingbeat. Despite possible inter-wingbeat variation, moths appear to use an unburst LEV with and without wind.

**Figure 8:**
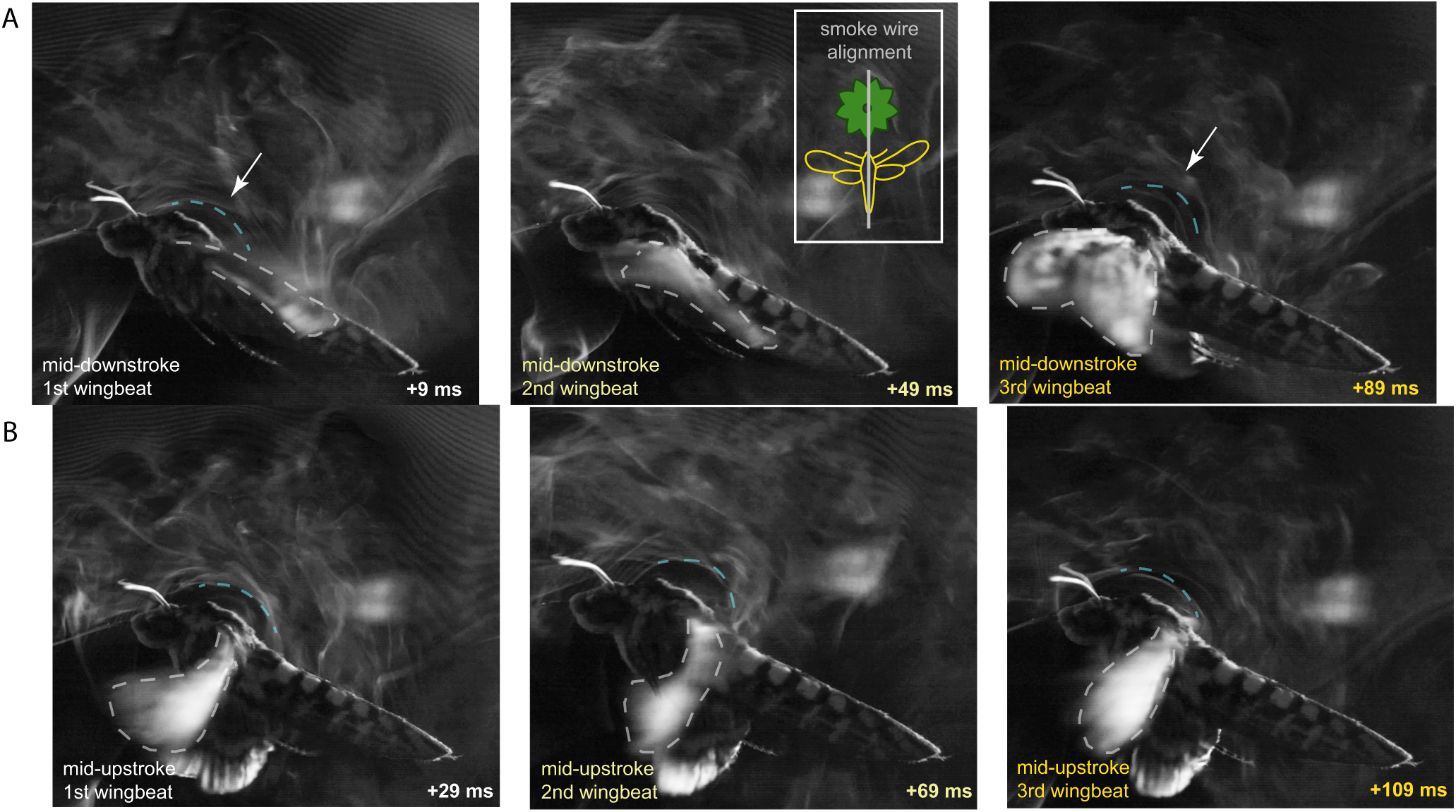
Smoke visualization of the leading edge vortex over the thorax (centerline). Flow is attached over the thorax at the late downstroke (for most wingstrokes), separates at stroke reversal, and then the LEV grows throughout the upstroke. Snapshots from three successive wing-beats for one moth show (A) a transient downstroke LEV over the thorax and (B) the persistent thorax LEV at mid-upstroke. Relative time throughout each wingbeat is shown based on the approximately 25 Hz (or 40 ms) wingbeat frequency (Fig. S2). When present (white arrow and blue dashed outline), the downstroke LEV is comparable in size to the upstroke LEV. The wing is outlined in gray. Inset shows smoke wire lateral alignment with the moth and flower.

## Discussion

### Wake interactions shift tracking performance within vortex shedding frequency range

Although performance declines, moths maintain near perfect tracking in the flower wake within the range of flower oscillations they encounter in nature. Outside of this range, moths face significant challenges from vortices shed in the flower wake. Gain overshoot, tracking error, and inertial power, all peak within 2-5 Hz and are higher than in still air (Figs. 3D,F and 6C,D,E). Hawkmoths (Ortega-Jimenez et al. 2013), bumblebees (Ravi et al. 2013), and fish (Liao et al. 2003*b,a*, Liao 2007, Maia et al. 2015) have all demonstrated an ability to stabilize perturbations at vortex shedding frequencies when maintaining a position. We found that shed vortices also impact active maneuvers, like flower tracking. Despite changes in underlying tracking dynamics, the flower wake does not lead to tracking failure. Aerodynamic interactions challenge tracking maneuvers, but moths still successfully feed and maintain comparable positional errors (Fig. S4).

Responses to the flower wake are consistent across all non-driving frequencies, regardless of whether the flower is moving or not. Wakes of oscillating cylinders exhibit increased vortex strength (Toebes 1969) and varied modes of vortex shedding depending on the frequency of oscillation (Griffin 1971, Williamson & Roshko 1988, Placzek et al. 2009). When the cylinder oscillates at the natural vortex shedding frequency of the still cylinder, a “lock-in” condition can be reached where the motion of the cylinder can drive vortex shedding away from the natural frequency (Koopmann 1967). Although the mean inertial power for moving and still flowers in wind differs slightly at non-driving frequencies outside of the vortex shedding range, the confidence intervals maintain overlap (Fig. 6D). The consistent overlap across non-driving frequencies in wind suggests that the vortex shedding in the flower wake occurs between 2-5 Hz whether the flower is moving or not (Fig. 6D). Separation between vortex shedding frequencies and flower motion frequencies may allow the moth to decouple tracking maneuvers and perturbation responses.

In nature, even wind speeds below 5 cm s^-1^ induce small amplitude oscillations (approximately 0.5 cm, peak-to-peak) in hawkmoth pollinated flowers (*Phlox*) and higher wind speeds result in larger lateral than longitudinal oscillations (Farina et al. 1994). Frequency analysis of multiple flower species showed that most hawkmoth-pollinated flowers oscillate below 5 Hz (Sprayberry & Suver 2011) with over 90% of flower power (power spectral density or oscillation energy at each frequency) contained below 1.7 Hz (Sponberg et al. 2015). In addition to oscillating within this low frequency band (0.1-1.7 Hz), we found that natural hawkmoth-pollinated flowers shed wake structures at these frequencies (Fig. 5). Because the frequencies of flower oscillation and vortex shedding overlap, natural flower wakes could pose challenges to tracking in nature. Alternatively, if the passive response to wind is in the same direction as tracking motion, then moths might exploit this phenomenon to “surf” on flower wakes.

### Nonlinear inertial power response indicates nonlinear tracking dynamics biased to low frequency motion

Tracking performance is reduced in the flower wake with a larger effect at vortex shedding frequencies. More motion than predicted is seen at the driving frequencies, especially at frequencies (2-5 Hz) where the flower sheds vortices (Figs. 4 and 6). Moths do not combine tracking and perturbation responses as a simple superposition. Hawkmoths have a high roll moment of inertia and are known to rely on passive damping mechanisms in response to roll perturbations (Hedrick et al. 2009, Cheng et al. 2011, Liu & Cheng 2017). Although the moment of inertia is lower around the yaw and pitch axes, similar passive mechanisms could be employed to stabilize position in the flower wake. Reliance on passive damping could explain the large overshoot and tracking error between 2-5 Hz (Figs. 3D,F), if the moth only actively tracks outside this frequency range. The addition of unsteady flow emphasizes that best tracking performance is biased to low frequencies matching natural flower oscillations.

Moths are perturbed by wind, but they may correct for this through feedback. Both experiments and numerical simulations showed that bumblebees in von Kármán streets respond to perturbations with passive, drag-based mechanisms (Hedrick et al. 2009, Ristroph et al. 2013), but must also use active flight control to maintain stability as wake perturbation effects increase over time (Ravi et al. 2016). In addition, we know the abdomen responds actively to visual motion stimuli to stabilize body pitch over multiple wingstrokes (Dyhr et al. 2013). Hawkmoths maneuvering in wind could adjust abdomen position either to increase drag and stabilize against perturbations or tilt their aerodynamic force vector without changing wing kinematics.

Measurements of energetic costs of tracking showed that maneuvering does increase COM inertial power, but the increase is small compared to the energy required for hovering alone (Sprayberry & Daniel 2007). At our freestream velocity, 0.7 ms^-1^, the moths are in slow forward flight, however wing kinematics and body pitch are comparable to hovering (Willmott et al. 1997). We only consider power due to lateral motion. Contributions from wing aerodynamic power could also change in wind and it is not clear how changes in wing kinematics would influence aerodynamic power. At higher wind speeds, hawkmoths pitch down and adopt a figure 8 wing path (Willmott & Ellington 1997*b*). Adding the unsteady wake on top of higher speed steady wind could exaggerate these adjustments. Although the flower wake could increase the total energy needed to hover, increased inertial power requirements for motion may not be accompanied by increased power output (or metabolic) demands. Rainbow trout adopt a Kármán gait to slalom though unsteady wakes and reduce costs of locomotion (Liao et al. 2003*a*). Moths could similarly slalom through the flower wake to track flower motion (Lehmann 2008, Lua et al. 2011, 2017). If so, they may reduce control against perturbations occurring at the driving frequencies, even if this results in increased overshoot (Fig. 3D). Energy from the flower wake disrupts tracking maneuvers, but vortices are thought to first interact with the moth aerodynamically. Then, the flower wake should first disrupt the LEV, which we can identify with smoke visualization of vortex bursting.

### LEV seen in steady air persists despite unsteady wake interactions

We see no evidence of vortex bursting due to interaction with the flower wake. Instead we observe a continuous LEV across the wings and thorax. In wind, the flower sheds vortices in multiple directions causing multidirectional flow separation at the leading edge of the wing, which may inhibit the ability of each wing to stabilize bound vortices, such as the LEV. To maintain LEV structure and size, energy must be dissipated or the vortex would continue to grow in size and strength until it is shed off of the wing into the wake of the insect. Interactions with the flower wake could alter spanwise flow (Birch & Dickinson 2001) and potentially induce vortex bursting. However, additional energy from the flower wake is successfully dissipated out of the LEV. The lack of vortex bursting suggests that the leading edge vortex is robust to interactions with vortex structures at the spatiotemporal scale of the robotic flower.

Stability of the LEV in wind could be due to the small size of vortex structures (Fig. 4), suggesting they may not be energetic enough to cause disruption. The comparable size of vortices in natural flower wakes (Fig. 5) also implies that vortex size is not a challenge to the LEV in nature. The temporal range of vortex shedding frequencies (2-5 Hz) lies well below the wingbeat frequency of the moth during hovering and tracking (approximately 25 Hz). We found that while interacting with wind at these frequencies moths still maintain the timescale of the wingbeat (Fig. S3). Therefore the timescale of LEV growth is also maintained in the unsteady flower wake. Nonetheless, wakes do affect tracking. Quantitative flow visualization of LEV strength and development throughout the wingstroke could reveal if lift forces are affected by interactions with wind although the LEV structure appears not to change.

### Counterintuitive reduced-order dynamics emerge in a more complex environment

Although LEV structure is qualitatively maintained, moths produce reduced-order dynamics for tracking in wind compared to still air. Interaction with the flower wake removes the bimodal response in both gain and phase within the range of vortex shedding frequencies (Fig. 3D,E). The wake could be considered a disturbance, but if the unsteady flow also alters force generation of the wings, then the underlying tracking dynamics have changed. In other words, the filter that transforms kinematics to forces is likely changed in the flower wake. Counterintuitively, this environmental filter simplifies tracking dynamics within 2-5 Hz. With lower frequency disturbances, such as the flower wake, moths may actively prioritize responses to lower frequencies and ignore the higher frequency flower motions. This is one way adding the flower wake could filter the tracking response of the moth. Electric fish vary opposing thrust and drag forces to remain stable in different flow speeds (Sefati et al. 2013). Passive responses to vortices in the flower wake could also counteract higher-order tracking maneuvers at certain frequencies, even if the moth does not deliberately control against these perturbations.

### Contradiction in wake impact on maneuvering and aerodynamics

Hawkmoths employ a stable leading edge vortex to produce lift despite changing how well they track flowers in still and windy conditions. Since vortices in the flower wake are relatively small, interactions between these structures and the LEV may result in vorticity or lift magnitude differences, but not cause the overall LEV to burst or change structure along the wingspan. LEVs have been classified for many different insects based on the qualitative structure of the vortex across the wingspan and a quantitative understanding of how energy is dissipated to maintain LEV stability (Bomphrey et al. 2005). Hawkmoths rely on a continuous, actively generated LEV in steady air (Willmott et al. 1997) and operate in the Reynolds number range of vortex bursting. Our results show that moths continue to use the same class of LEV with no obvious evidence of vortex bursting.

While smoke visualization can identify LEV bursting, it cannot quantify changes in spanwise flow, either through the vortex core or towards the trailing-edge of the wing. We conclude that any changes in spanwise flow due to the flower wake are not large enough to burst the LEV or change its class.

### Consequences for flight control

Freely behaving animals rely on feedback systems to control locomotion and encode information about both the environment and their state within that environment. Fish swimming through von Kármán cylinder wakes largely maintain position through visual feedback (Liao 2007). The foraging task studied here was previously shown to rely on redundant visual and mechanosensory (from the proboscis-nectary interaction) pathways (Roth et al. 2016). The addition of mechanosensory feedback may help moths stabilize in wakes even if their vision is compromised. *M. sexta* also has a longer proboscis than some other hawkmoths, *D. elpenor* or *M. stellatarum* (Haverkamp et al. *2016),* which has been proposed to increase mechanosensory feedback during tracking since more of the proboscis is in contact with the nectary (Stöckl et al. 2017). A longer proboscis could also cause the moth to interact with vortices that develop further downstream from the flower, requiring the LEV to also be stable to the wakes of a wide range of flowers from *Petunia* to larger *Datura* (Fig. 5).

As they maneuver in nature, insects may need to encode both the environment around them and the forces they are able to produce through interactions with that environment. Campaniform sensilla present along the dorsal and ventral side of the hawkmoth forewing are sensitive to inertial forces on a timescale 80x smaller than the wingbeat period (Dickerson et al. 2014, Pratt et al. 2017). In unsteady flow, aerodynamic interactions change local flow along the wing and lead to LEV bursting. Aerodynamic forces are not thought to influence overall wing motions, but they may locally strain the wing if changes in flow are large. If the boundary layer is disrupted, wind may also deflect hair sensilla on thorax. Encoding local strains could be a mechanosensory feedback mechanism for LEV stability.

Although performance suffers, the forces and torques fundamental to successful flower tracking at natural flower oscillation frequencies are maintained in wind. Features of the environment are sensed and integrated by multiple neural pathways to achieve a desired motion. Variation in the environment then shifts behavior away from ideal locomotion, but not far enough to cause failure. Animals in nature may depend on sensing subtle changes in force-generating mechanisms, such as the LEV, to balance body maneuvers and lift production in an unsteady environment.

## Acknowledgements

We thank Steven Chandler for his assistance in setting up and maintaining the animals and experimental apparatus and many thoughtful discussions of the frequency domain. We also thank Isabel Veith for preparing supplementary information about the wind tunnel specifications.

## Competing interests

The authors declare no competing or financial interests.

**Author Contributions**

Conceptualization, methodology, resources, writing – review and editing, and funding acquisition: M.G.M. and S.N.S.; Software, validation, formal analysis, investigation, data curation, visualization, and writing – original draft and preparation: M.G.M.; Supervision: S.N.S.

## Supplementary Information

### Wind tunnel characteristics

The Eiffel-type wind tunnel used in this experiment is located at Georgia Institute of Technology (Atlanta, GA, USA). It was designed, constructed, and tested by Engineering Laboratory Design, Incorporated (Lake City, MN, USA). It is a low speed, open circuit wind tunnel situated in a room with dimensions 11.3 x 24.88 x 2.96 m (length x width x height). Air speeds range from 0.25-10 m s^-1^, with velocity variation less than ±2% of the mean free stream velocity. Based on the maximum free stream velocity, streamwise turbulence should not exceed 0.5%. Air is drawn into the elliptical inlet and passes through a honeycomb mesh to condition the flow, which is subsequently contracted and accelerated through to the test section. To regain static pressure, the air then passes through a diffuser before traveling through the fan and back into the room. The entire wind tunnel is supported by structural steel frames positioned on fitted leveling pads. To prevent vibrations of the fan and room from interfering with the air flow, the wind tunnel sections and supporting frames are isolated with rubber-shear-in mounts and flexible coupling.

### Flow Ducts and Straighteners

The ducts are a composite of fiberglass and reinforced plastic with a molded balsa wood core. The inlet of the settling chamber and flow ducts has 1.61 m lateral clearance to the walls and 0.23 m vertical clearance to the ceiling. Prior to the contraction section, air passes first through a settling chamber and then through a tensioned, hexagonal-cell aluminum honeycomb grid. Before air enters the test section, it flows through a duct fitted with flow straighteners to maximize laminar flow into the test section. The contraction section has a 6.25:1 area ratio with a symmetric cross-section. The inlet and exit areas of this section have static pressure taps.

### Test Section

Moths were flown in the test section which is joined to the diffuser and contraction sections via aluminum angle flanges. The test section is 1.37 m from the walls of the room and can (optionally) be divided into two separate, but continuous sections. The primary section (100 cm) is double the length of the secondary test section (50 cm). Test sections are accessible by way of portholes in the panels. The floor, ceiling, and sidewalls of the primary section are made of 13 mm thick soda lime glass. Sidewalls are secured with toggle clamps and can be removed and replaced with custom panels depending on experimental needs. On the operator side of the test section, the sidewall panel has a pneumatic opening door for easy access to the test section. The ceiling, floor, and sidewalls of the secondary section are made of 19.1 mm thick, clear, GM grade acrylic. There are 34M stainless steel, high porosity (60%), tensioned catch screens in place on either side of the test section to prevent the moths from flying into other components of the wind tunnel.

### Diffuser

Air flows out from the test section into the diffuser which expands with a total angle of 4.6^*o*^ and is separated from the fan by an air gap. This air gap (13.9 cm in width and 2.43 m circular cross-section diameter) acts as a vibration isolator. The diffuser contains a highly porous perforated plate to decelerate the flow before it exits into the room and protects the fan.

### Fan and Fan Motors

Airflow is produced by an inline, centrifugal belt driven fan with a 3.0 HP, ODP 208/230 VAC induction motor. The fan is 2.13 m from the door and controlled by a transistor inverter variable frequency speed controller (208-230 VAC/60 Hz/30 Amp). The fan is equipped with a fusible disconnect to protect the motor and controller. A remote control operator station is located downstream of the test section, near the upstream end of the diffuser.

### Frequency response and tracking performance

First, we Fourier transform the individual time series data for each trial,

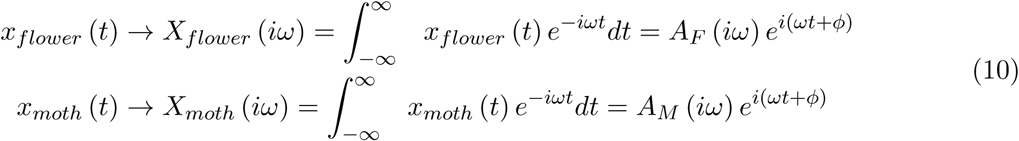

We then use the complex ratio of the Fourier transformed moth and flower motion to define gain (*G*) as the absolute value of the complex ratio

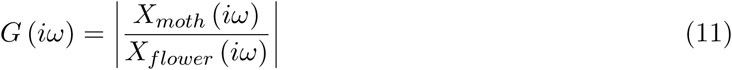

and phase (*ϕ*) as the angle between the real and imaginary parts of the complex response ratio

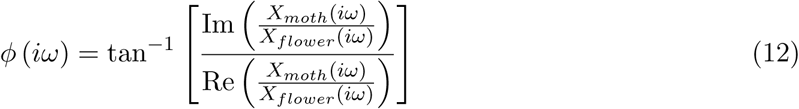

Respectively, these quantities represent the relative position and timing differences between flower and moth. This frequency response can be represented in the (polar) complex plane by ordered pairs (*G, ϕ*) where gain corresponds to the radial distance and phase to the angle measured counterclockwise from the horizontal. In this representation, perfect flower tracking occurs at (1,0). To explore how tracking performance (measured here using gain and phase) varies with frequency, we plot gain and phase separately, but both are necessary to describe the behavior of the system across all frequency bands.

## Supplementary Figures

**Figure S1:**
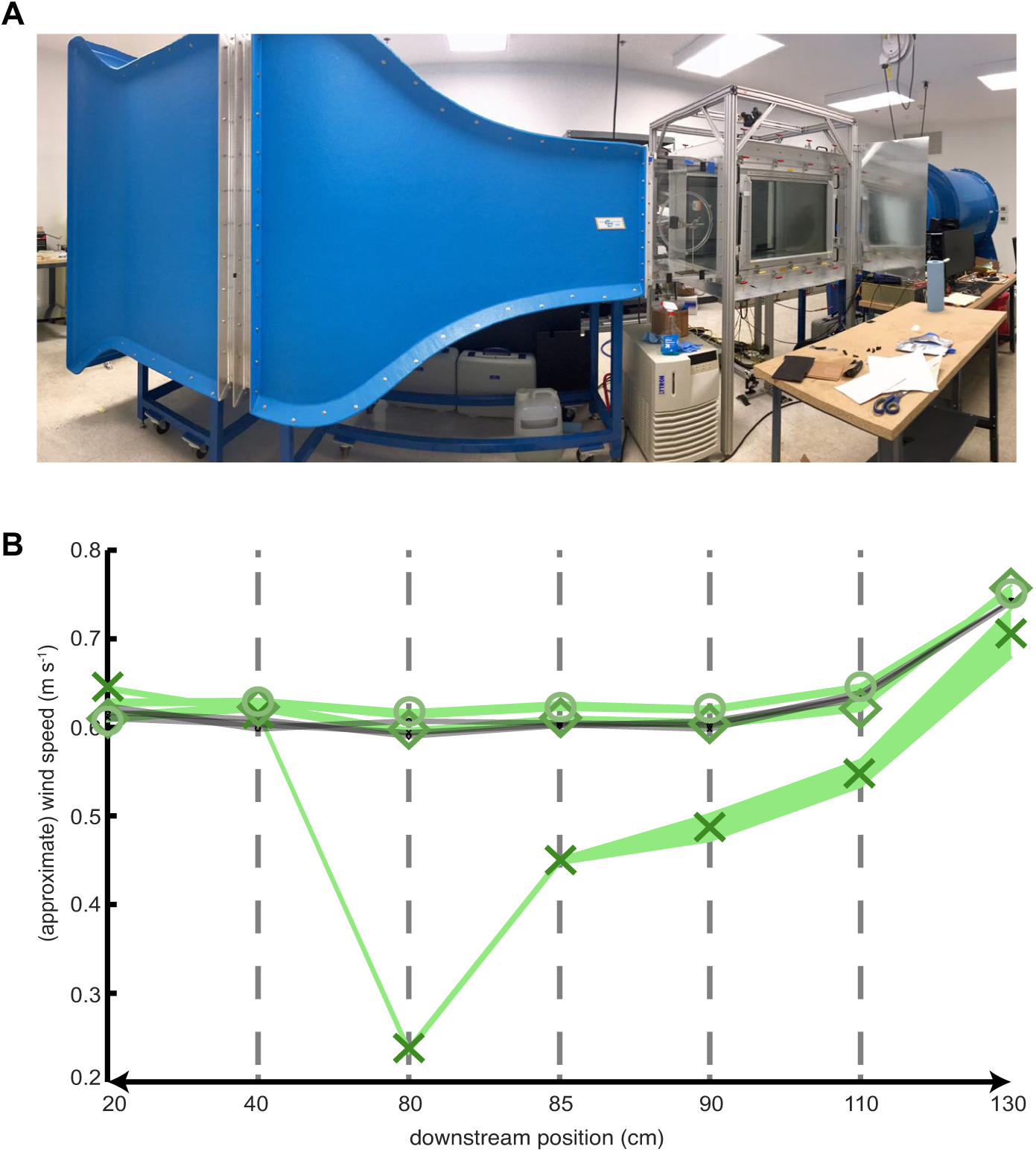
Wind tunnel and flow profile. (A) Picture of wind tunnel setup. (B) Wind speed measurements v. downstream position with (green) and without (black) the flower present. Measurements were taken at three lateral (x-direction) positions spaced 4 cm apart (shown in Fig. 2A), but only flow along the centerline showed changes due to the flower. Symbols for the flower wake are enlarged for clarity. Downstream positions are: (1) 20 cm downstream of front mesh, (2) 40 cm downstream of front mesh, (3) expected moth position (80 cm downstream), (4) 85 cm downstream of front mesh, (5) 90 cm downstream of front mesh, (6) 110 cm downstream of front mesh, and (7) 130 cm downstream of front mesh (20 cm upstream of back mesh). Without the flower present, the flow speed profile was uniform across the full length of the working section with variation ±0.01 ms^-1^ (black data points). With the flower, all flow variation occurred downstream, along the centerline (green x-symbol). In this region, flow speed increases from approximately 0.2 ms^-1^ at the expected moth position (approximately 80 cm downstream), to 0.6 ms^-1^ at 110 cm downstream with velocity variation ±0.02 ms^-1^. At 130 cm downstream wake effects of the flower were dissipated and the flow speed once again matched the 0.7 ms^-1^ free-stream velocity. Flow upstream of the flower was uniform and consistent with velocity measurements made without the flower present.

**Figure S2:**
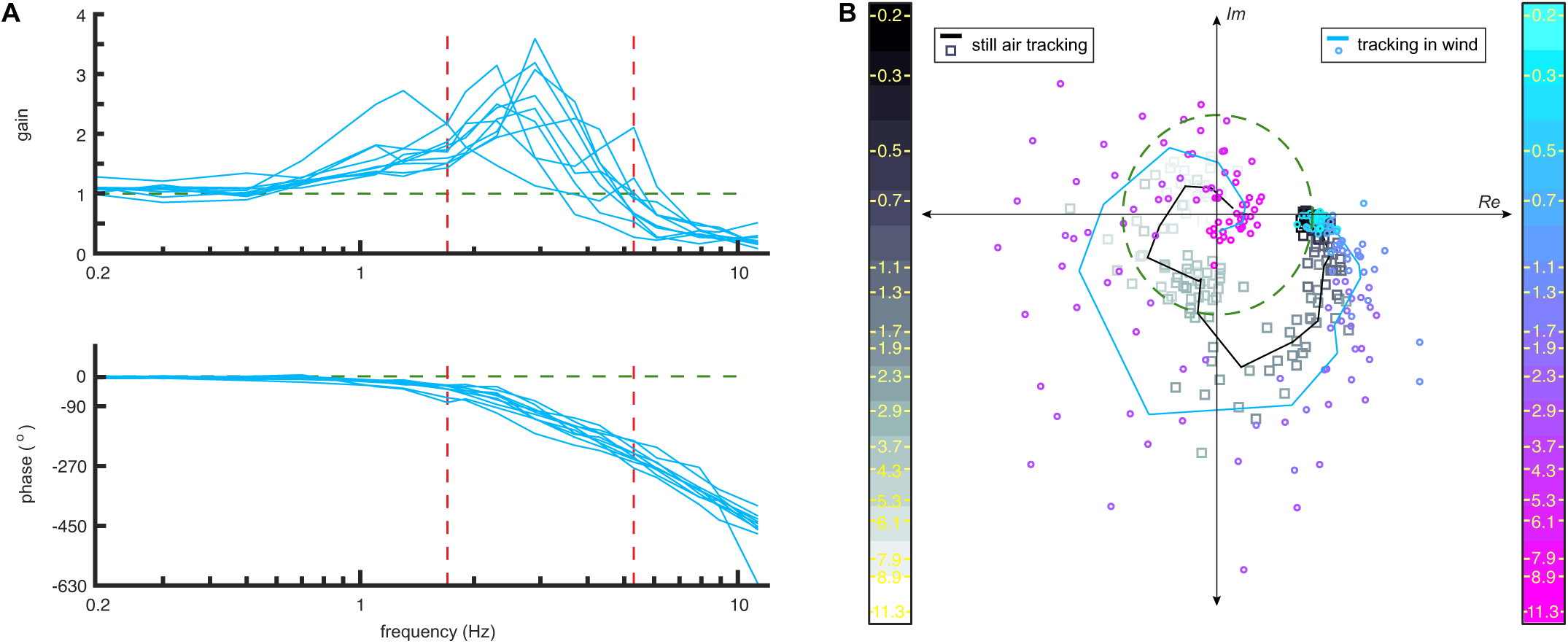
Individual bode and average complex tracking response with and without wind. (A) Gain and phase of all 10 individual moths for tracking in wind. The region of vortex shedding frequencies is bounded between 1.7 Hz and 5.3 Hz (red lines). (B) Complex tracking response with and without wind. Perfect tracking corresponds to (1,0) on the green unit circle. Gain and phase are represented in terms of radius and angle around the circle, respectively. Data points correspond to individuals and solid lines represent the mean. Color bars identify how tracking varies across all driving frequencies in still air (greyscale squares) and wind (cool scale, circles). All still air data (black line, grayscale squares) previously collected in Sponberg et al. (2015).

**Figure S3:**
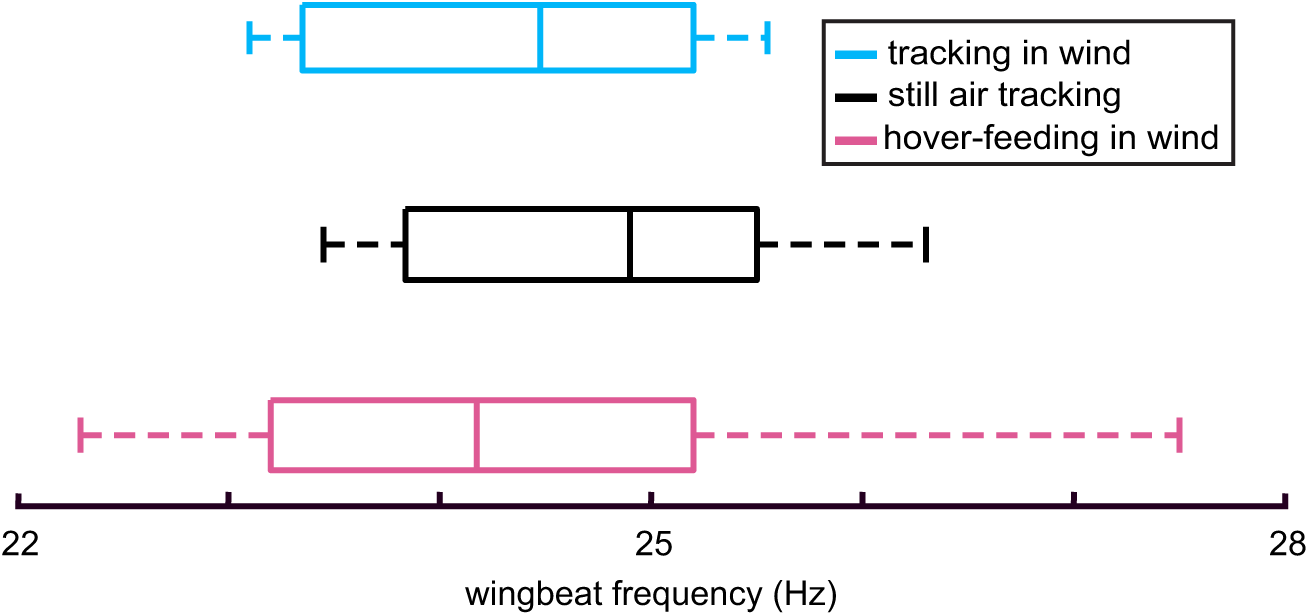
Wingbeat frequency shows relatively little change across experimental conditions. Boxplots of wingbeat frequency for tracking in wind (blue), still air tracking (black), and hover-feeding in wind (pink). Median wingbeat frequency varies by less than 1 Hz between the three conditions. Whiskers correspond to the most extreme data points and the boundaries of the box denote the 25th and 75th data percentiles.

**Figure S4:**
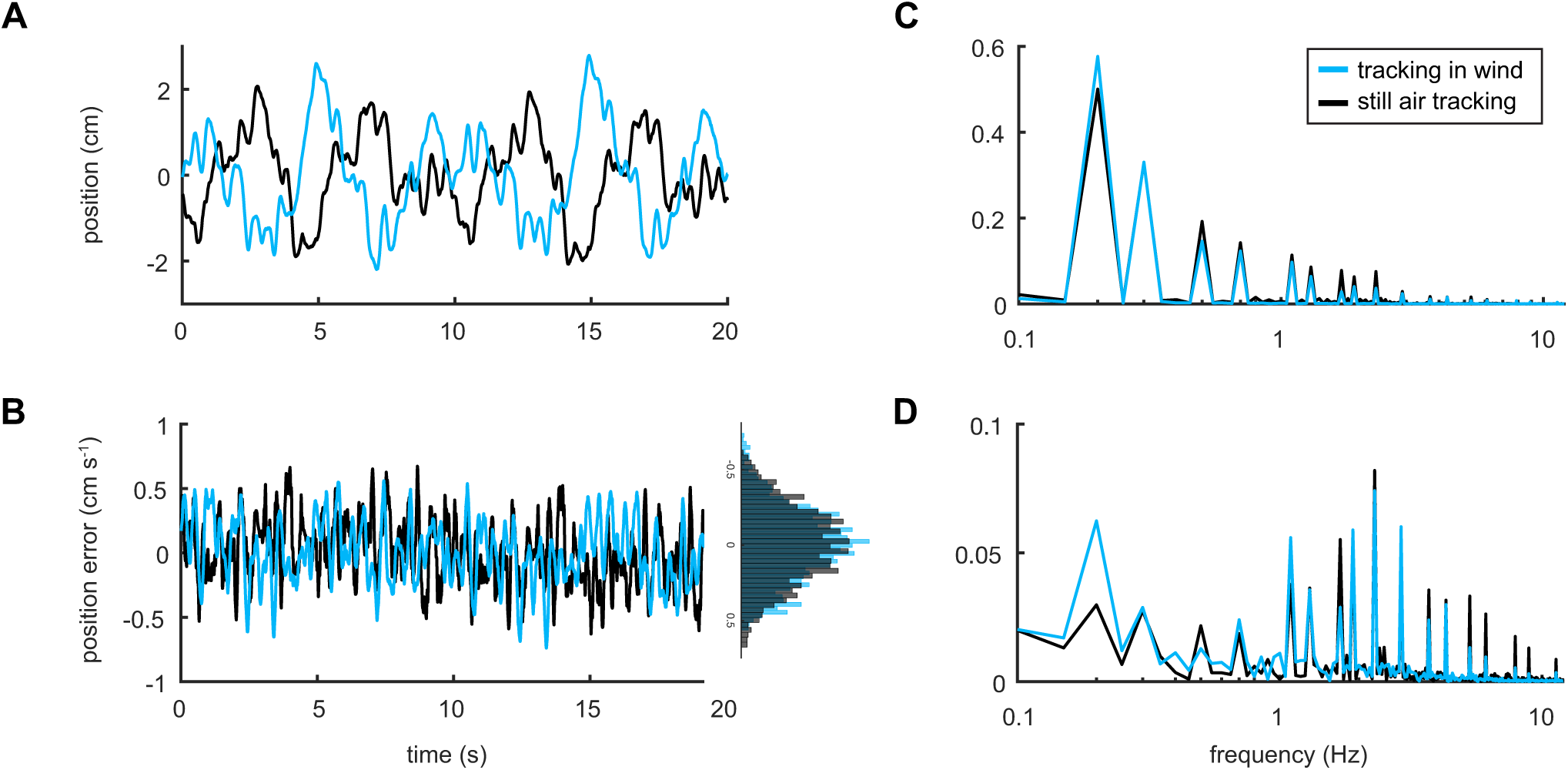
Position error in the flower wake. All still air data (black) previously collected in Sponberg et al. (2015). (A) Averaged time series data for tracking with and without wind. (B) Position error with and without wind. Differences in the time traces are emphasized with the inset histogram. In wind, position error increases and moths spend more time further from the flower. (C) Amplitude (position) in the frequency domain (after Fourier transform of data from A). Peaks correspond to prescribed flower driving frequencies. (D) Position error in the frequency domain (after Fourier transform of data from B). Tracking is dominated by low frequency oscillations due to the stimulus design. In wind, position error increases across low frequencies, but decreases at high frequencies. Across all frequencies tracking performance is decreased and higher frequency oscillations are removed.

**Figure S5:**
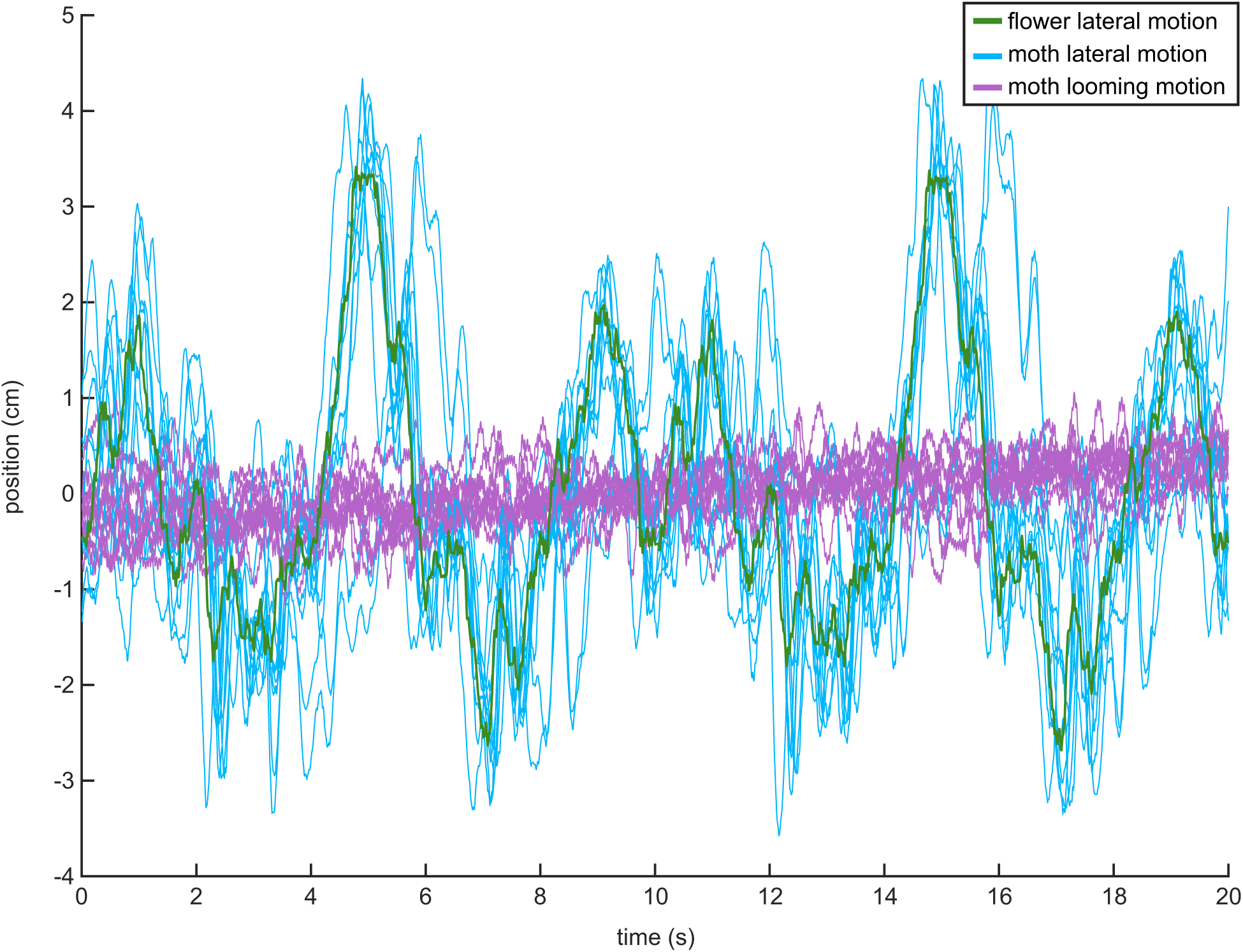
Lateral oscillations dominate tracking motion. Raw time series data for individual trials of tracking in wind. The flower motion (green) is bounded between ±3 cm in the x-direction, but the moth (blue) overshoots the flower sometimes reaching 4 cm lateral displacements. Off-axis oscillations in the looming, or y-direction, are small compared to lateral motion. Moths maintain position ±1 cm downstream of the flower.

## Supplementary Videos

Movie S1. **Smoke visualization video of the robotic flower wake.** Smoke plane is aligned with the center of the flower and playback is at 30 fps (recorded at 125 fps). Vortices in the flower wake are shed within the frequency range 2-5 Hz. The flower wake is best described by a range of frequencies due to the complex shape of the flower face and nectary combined. Global level and contrast adjustments were performed in iMovie to better visualize the smoke lines.

Movie S2. **Smoke visualization video of the leading-edge vortex (LEV) at mid-wing.** Smoke plane is aligned with mid-wing on the moth (approximately 3-4 cm off center) and playback is at 30 fps (recorded at 125 fps). Although the wings are interacting with vortices shed off of the flower, the LEV remains bound to the wing throughout the downstroke and is best seen during mid-downstroke. Global level and contrast adjustments were performed in iMovie to better visualize the smoke lines.

Movie S3. **Smoke visualization video of the leading-edge vortex (LEV) over the thorax.** Smoke plane is aligned with the center of the flower and playback is at 30 fps (recorded at 125 fps). Although the wings are interacting with vortices shed off of the flower, the LEV remains bound to the wing throughout the downstroke and is best seen during mid-downstroke. Global level and contrast adjustments were performed in iMovie to better visualize the smoke lines.

